## Supplemental File for "High-level production of nervonic acid in the oleaginous yeast *Yarrowia lipolytica* by systematic metabolic engineering"

***Corresponding author:**

#### Supplementary Tables

##### Table S1. The *Y. lipolytica* strains used in this study

| Strain | Genotype | Source |
|---|---|---|
| po1g | *MATa, leu2-270, ura3-302::URA3, xpr2-3* | Yeastern |
| po1g-G3 | po1g, *ACC1, DGA1, SCD, ΔKu70* | (Li et al., 2020) |
| YL-AtADS2 | po1g, *AtADS2* | This work |
| YL-CgKCS | po1g, *CgKCS* | This work |
| YL-erCgKCS | po1g, *ER-CgKCS* | This work |
| YL-mtCgKCS | po1g, *MT-CgKCS* | This work |
| YL-peCgKCS | po1g, *PE-CgKCS* | This work |
| YL-erpeCgKCS | po1g, *ER-CgKCS*, *PE-CgKCS* | This work |
| YL-AtFAE1 | po1g, *AtFAE1* | This work |
| YL-BtFAE1 | po1g, *BtFAE1* | This work |
| YL-3FAEs | po1g, *AtFAE1*, *BtFAE1*, *CgKCS* | This work |
| YL-gfpKCS | po1g, *CgKCS-sfGFP* | This work |
| YL-gELOVE6 | po1g, *gELOVE6* | This work |
| YL-MaOLE2 | po1g, *MaOLE2* | This work |
| YL-3CgK | po1g, (*ER-CgKCS, MT-CgKCS, PE-CgKCS*)*2 | This work |
| YL-3CgKE | YL-3CgK, *gELOVE6* | This work |
| YL-3CgKEM | YL-3CgKE, *gELOVE6*, *MaOLE2* | This work |
| YL-ΔFAD2 | po1g, *FAD2::URA3* | This work |
| YLVL3 | po1g-G3, *ER-CgKCS*, *MT-CgKCS*, *PE-CgKCS* | This work |
| YLVL6 | YLVL3, *ER-CgKCS, MT-CgKCS, PE-CgKCS* | This work |
| YLVL7 | YLVL6, rDNA::(*CgKCS-gElovl6-MaOLE2*) | This work |
| YLVL8 | YLVL7, D17::(*CgKCS-gElovl6-MaOLE2*) | This work |
| YLVL10 | YLVL8, *FAD2*::*CgKCS**2 | This work |
| YLNA1 | po1g-G3, rDNA::*CgKCS* | This work |
| YLNA3 | YLNA1, *FAD2*::*CgKCS**2 | This work |
| YLNA5 | YLNA3, *TGL4*::*CgKCS**2 | This work |
| YLNA6 | YLNA5, *GSY1*::*CgKCS* | This work |

| | | |
|---|---|---|
| YLNA7 | YLNA6, *SNF1*::*CgKCS* | This work |
| YLNA8 | YLNA7, *D17*::(*CgKCS-MaOLE2*) | This work |
| YLNA9 | YLNA8, *PEX10*::(*YlINO2*) | This work |
| YLNA10 | YLNA8, *PEX10*::(*MoGPAT-88*) | This work |

Note: ER, endoplasmic reticulum; MT, mitochondria; PE, peroxisome.

**Table S2. Plasmids used in this study**

| Plasmid | Genotype | Source |
|---|---|---|
| pYLEX1 | p*hp4d*, *LEU2* | Yeastern |
| pYLU | pYLEX1, *LEU2::URA3* | This work |
| pYL-AtADS2 | pYLU, *TEFin-AtADS2* | This work |
| pYL-CgKCS | pYLU, *TEFin-CgKCS* | This work |
| pYL-erCgKCS | pYLU, *TEFin-ER-CgKCS* | This work |
| pYL-mtCgKCS | pYLU, *TEFin-MT-CgKCS* | This work |
| pYL-peCgKCS | pYLU, *TEFin-PE-CgKCS* | This work |
| pYL-AtFAE1 | pYLEX1, *TEFin-AtFAE1* | This work |
| pYL-erAtFAE1 | pYLEX1, *TEFin-ER-AtFAE1* | This work |
| pYL-peAtFAE1 | pYLEX1, *TEFin-PE-AtFAE1* | This work |
| pYL-BtFAE1 | pYLEX1, *TEFin-BtFAE1* | This work |
| pYL-erBtFAE1 | pYLEX1, *TEFin-ER-BtFAE1* | This work |
| pYL-peBtFAE1 | pYLEX1, *TEFin-PE-BtFAE1* | This work |
| pYL-gfpKCS | pYL-CgKCS, *TEFin-CgKCS-sfGFP* | This work |
| pYL-gELOVE6 | pYLU, *YAT1-gELOVE6* | This work |
| pYL-MaOLE2 | pYLU, *YAT1-MaOLE2* | This work |
| pYL-ΔFAD2 | pYLU, *FAD2up-URA3-FAD2dn* | This work |
| pYL-2KCS | pYL-erCgKCS, *TEFin-MT-CgKCS* | This work |
| pYL-3sKCS | pYL-2KCS, *TEFin-PE-CgKCS* | This work |
| pYL-3grDNA | pYL-CgKCS,*rDNAup-CgKCS-gElovl6-MaOLE2-rDNAdn* | This work |
| pYL-3gD17 | pYL-3grDNA,*D17up-CgKCS-gElovl6-MaOLE2-D17dn* | This work |
| pYL-2gFAD2 | pYL-ΔFAD2, *(TEFin-CgKCS)* *2 | This work |
| pYL-rDNA-CgKCS | pYL-3grDNA, *rDNAup-CgKCS -rDNAdn* | This work |
| pYL-2gTGL4 | pYL-2gFAD2, *TGL4up-CgKCS-CgKCS-TGL4dn* | This work |
| pYL-GSY1-CgKCS | pYL-rDNA-CgKCS, *GSY1up-CgKCS-GSY1dn* | This work |
| pYL-SNF1-CgKCS | pYL-rDNA-CgKCS, *SNF1up-CgKCS-SNF1dn* | This work |

| | | |
|---|---|---|
| pYL-D17-gMaolE2 | pYL-rDNA-CgKCS, *D17up-CgKCS-MaolE2-D17dn* | This work |
| pYL-PEX10-YlINO2 | pYL-rDNA-CgKCS, *PEX10up-YlINO2-PEX10dn* | This work |
| pYL-PEX10-YlINO4 | pYL-rDNA-CgKCS, *PEX10up-YlINO4-PEX10dn* | This work |
| pYL-PEX10-Mo-35 | pYL-rDNA-CgKCS, *PEX10up-MoDGAT-35-PEX10dn* | This work |
| pYL-PEX10-Mo-49 | pYL-rDNA-CgKCS, *PEX10up- MoDGAT-49-PEX10dn* | This work |
| pYL-PEX10-Mo-88 | pYL-rDNA-CgKCS, *PEX10up-MoGPAT-88-PEX10dn* | This work |
| pYL-PEX10-Mo-90 | pYL-rDNA-CgKCS, *PEX10up-MoGPAT-90-PEX10dn* | This work |

### Table S3. Primers used for construction of plasmids

| Name | Sequence (5' to 3') | PCR template | Target plasmid |
|---|---|---|---|
| pYL-F | ctgcggttagtactgcaaaaagtgctg | pYLU | pYL-AtADS2 |
| pYL-R | ccccacgttgccggtcttgc | | |
| ADS2-F | actttttgcagtactaaccgcagtctgtgacctctaccgtggag | Synthesized *AtADS2* gene | |
| ADS2-R | gcaagaccggcaacgtggggtcatctcacgatggccattc | | |
| CgKCS-F | actttttgcagtactaaccgcagacctctatcaacgtgaagctg | Synthesized *CgKCS* gene | pYL-CgKCS |
| CgKCS-R | gcaagaccggcaacgtggggttaagatcggccgttttggg | | |
| CgKer-F | cggccgatcttaaggacgagctgtaaccccacgttgccggtcttg | pYL-CgKCS | pYL-erCgKCS |
| CgKer-R | cgtggggttacagctcgtccttagatcggccgttttgggctc | | |
| CgKpe-F | cggccgatcttccaagctgtaaccccacgttgccggtcttg | pYL-CgKCS | pYL-peCgKCS |
| CgKpe-R | cgtggggttacagcttggaagatcggccgttttgggctc | | |
| pCgKmt-F | aagcagatatctagagctacac | p-eGFP-mt | pYL-mtCgKCS |
| pCgKmt-R | gtatctacacgcgtgctatgcatctgg | | |
| CgKmt-F | catagcacgcgtgtagatacttaagatcggccgttttggg | pYL-CgKCS | |
| CgKmt-R | gtagctctagatatctgcttatgacctctatcaacgtgaag | | |
| pYL1-F | gtatctacacgcgtgctatg | pYLEX1 | pYL-AtFAE1 |
| pYL1-R | tgttgatgtgtgtttaattcaag | | |
| AtE1-F | catagcacgcgtgtagatacttaagatcggccgttctgcac | Synthesized *AtFAE1* gene | |
| AtE1-R | gaattaaacacacatcaacaatgacctctgtgaacgtgaag | | |
| AtE1er-F | tagatacttacagctcgtccttagatcggccgttctgcacgt | pYL-AtFAE1 | pYL-erAtFAE1 |
| AtE1er-R | cggccgatctaaggacgagctgtaagtatctacacgcgtgctatg | | |
| AtE1pe-F | tagatacttacagcttggaagatcggccgttctgcacgt | pYL-AtFAE1 | pYL-peAtFAE1 |
| AtE1pe-R | cggccgatcttccaagctgtaagtatctacacgcgtgctatg | | |
| pYL2-F | ctgcggttagtactgcaaaaagtgctg | pYL-CgKCS | pYL-BtFAE1 |
| pYL2-R | ccccacgttgccggtcttgc | | |
| BtE1-F | actttttgcagtactaaccgcagacctctgtgaacgtgaag | Synthesized *BtFAE1* gene | |
| BtE1-R | gcaagaccggcaacgtggggttaagatcggccgttctg | | |
| BtE1er-F | tagatacttacagctcgtccttagatcggccgttctgcacgt | pYL-BtFAE1 | pYL-erBtFAE1 |
| BtE1er-R | cgtggggttacagctcgtccttagatcggccgttctgcactc | | |
| BtE1pe-F | tagatacttacagcttggaagatcggccgttctgcacgt | pYL-BtFAE1 | pYL-peBtFAE1 |
| BtE1pe-R | cgtggggttacagcttggaagatcggccgttctgcactc | | |
| pYL3-F | taaccccacgttgccggtc | pYL-CgKCS | pYL-gfpKCS |
| pYL3-R | agatcggccgttttgggc | | |
| sfG-F | gagcccaaaacggccgatctgcagtgtcttgcgtctctg | Synthesized *sfGFP* gene | |
| sfG-R | agaccggcaacgtggggttacttgtaaagctcatccatgcc | | |
| pYL4-F | ctgcggttagtactgcaaaaagtgctg | pYL-CgKCS | pYL-TEF-gELOVE6 (TEF1in promoter) |
| pYL4-R | ccccacgttgccggtcttgc | | |
| gEL6-F | actttttgcagtactaaccgcagtctgtgctgaccctgcagga | Synthesized *gELOVL6* gene | |
| gEL6-R | gcaagaccggcaacgtggggttagtcggccttggtggcct | | |
| pYL5-F | ctgcggttagtactgcaaaaagtgctg | pYL-CgKCS | pYL-TEF-MaOLE2 (TEF1in promoter) |
| pYL5-R | ccccacgttgccggtcttgc | | |
| MaO2-F | actttttgcagtactaaccgcaggccaccccccctgcctcctac | Synthesized *MaOLE2* gene | |
| MaO2-R | gcaagaccggcaacgtggggttactcctccttagagtggtcgccg | | |
| pE6-F | taattcacaaatgtctgtgctgaccctg | pYL-TEF-gELOVE6 | pYL-gELOVE6 (Yat1 promoter) |
| pE6-R | atggcgaagggtcgaccgatgcccttgag | | |
| Ye6-F | atcggtcgacccttcgccatgcaatttg | *Y. lipolytica* gDNA | |
| Ye6-R | gcacagacatttgtgaattagggtggtg | | |
| pMa-F | taattcacaaatggccacccccctgcctcc | pYL-TEF-MaOLE2 | pYL-MaOLE2 |

| Name | Sequence (5’ to 3’) | PCR template | Target plasmid |
|---|---|---|---|
| pMa-R | atggcgaagggtcgaccgatgcccttgagagcc | | (Yat1 promoter) |
| Ymao-F | atcggtcgacccttcgccatgcaatttg | *Y. lipolytica* gDNA | |
| Ymao-R | gggtggccatttgtgaattagggtggtg | | |
| pF2up-F | aattccgtcgtcgcctgagtc | pYL-CgKCS | pYL-FAD2up |
| pF2up-R | gtcgaccgatgcccttgagag | | |
| F2up-F | tctcaagggcatcggtcgacctacggcacgataaagatgg | *Y. lipolytica* gDNA | |
| F2up-R | actcaggcgacgacggaattagctagaaatgttatttgattgtg | | |
| pF2dn-F | gatctcggcagtctctcggatg | pYL-ΔFAD2-up | pYL-ΔFAD2 |
| pF2dn-R | tgttcggaaatcaacggatg | | |
| F2dn-F | catccgttgatttccgaacattctatggtcggtctgttgg | *Y. lipolytica* gDNA | |
| F2dn-R | tccgagagactgccgagatcttacaatgggaccgtgcttg | | |
| pCgER-F | aattccgtcgtcgcctgagtc | pYL-erCgKCS | pYL-2KCS |
| pCgER-R | ggacacgggcatctcacttgcatatgtatg | | |
| Cgmt-F | caagtgagatgcccgtgtccctaccttgctcgaatgacttattgtg | pYL-mtCgKCS | |
| Cgmt-R | actcaggcgacgacggaattcgcagtaggatgtcctgcac | | |
| pCpe-F | agagaccgggttggcggcgc | pYL-peCgKCS | pYL-3sKCS |
| pCpe-R | gtcgaccgatgcccttgagag | | |
| 3Csig-F | aaggctctcaagggcatcggtcgacagagaccgggttggcggcgc | pYL-2KCS | |
| 3Csig-R | caaatgcgccgccaacccggtctctcgcagtaggatgtcctgcacggg | | |
| rdc-1F | gtgttacctacaggacatcctactgcgag | pYL-CgKCS | pYL-rDNA-CgKCS |
| rdc-1R | atctgcactatgttcggaaatcaacggatg | | |
| 26yup-F | tggcggccgcttcacctccttcatccgag | *Y. lipolytica* gDNA | |
| 26yup-R | ggatgtcctgtaggtaacacgcttaagcg | | |
| rdc-2F | tcttcctatcgcggccgcgatctcggcagtctctcg | pYL-CgKCS | |
| rdc-2R | aggaggtgaagcggccgccagagcagattgtactgagag | | |
| 26ydn-F | tttccgaacatagtgcagatcttggtgg | *Y. lipolytica* gDNA | |
| 26ydn-R | tcgcggccgcgataggaagagccgacatc | | |
| pYgE-F | gcccgtgtccgctctcaagggcatcggtc | pYL-gELOVE6 | pYL-gELovl6-MaOLE2 |
| pYgE-R | atggcgaaggcttcaacccagtcagctcc | | |
| MoY-F | tgggttgaagccttcgccatgcaatttgtc | pYL-MaOLE2 | |
| MoY-R | ccttgagagcggacacgggcatctcacttg | | |
| prCg-F | gcccgtgtcccaggacatcctactgcgag | pYL-rDNA-CgKCS | pYL-3grDNA |
| prCg_R | atggcgaaggtaggtaacacgcttaagcg | | |
| gEMa-F | gtgttacctaccttcgccatgcaatttgtc | pYL-gELovl6-MaOLE2 | |
| gEMa-R | ggatgtcctgggacacgggcatctcacttg | | |
| pD17-1F | tgcagaaactcgggatggcgcttaagcgtg | pYL-3grDNA | pYL-3gD17 |
| pD17-1R | gagacttcgctctgttcggaaatcaacggatgct | | |
| 17dn-f | gatttccgaacagagcgaagtctcgctggac | *Y. lipolytica* gDNA | |
| 17dn-r | tcgaaggatcaatcctgtccaagctagtcttctatc | | |
| pD17-2F | gcttggacaggattgatccttcgatgtcgg | pYL-3grDNA | |
| pD17-2R | cagacactgcgtgtaatctcggatgaaggag | | |
| 17up-f | atccgagattacacgcagtgtctgaatgttc | *Y. lipolytica* gDNA | |
| 17up-r | taagcgccatcccgagtttctgcagcatatc | | |
| pFAD-1F | gatgtcatggaagaaggagctgactgggttg | pYL-2KCS | pYL-2gFAD |
| pFAD-1R | aacgagttcacttgttcggaaatcaacggatg | | |

| Name | Sequence (5' to 3') | PCR template | Target plasmid |
|---|---|---|---|
| FADdn-F | gatttccgaacaagtgaactcgttgccaaag | *Y. lipolytica* gDNA | |
| FADdn-R | tcgaaggatcaatgtgcaagagtgtgtacg | | |
| pFAD-2F | cactcttgcacattgatccttcgatgtcgg | pYL-2KCS | |
| pFAD-2R | acaaaccctgccgtaatctcggatgaaggag | | |
| FADup-F | atccgagattacggcagggtttgttggtgttaatg | *Y. lipolytica* gDNA | |
| FADup-R | gtcagctccttcttccatgacatcatcgagacc | | |
| pTGL4-1F | catcgaagactagaaggagctgactgggttg | pYL-2gFAD | pYL-2gTGL4 |
| pTGL4-1R | tgcgaatgcttatgttcggaaatcaacggatg | | |
| TGL4dn-F | gatttccgaacataagcattcgcagtcgtc | *Y. lipolytica* gDNA | |
| TGL4dn-R | tcgaaggatcaactatgtaattgaccagaaacag | | |
| pTGL4-2F | tcaattacatagttgatccttcgatgtcgg | pYL-2gFAD | |
| pTGL4-2R | cataagtagctagtaatctcggatgaaggag | | |
| TGL4up-F | atccgagattactagctacttatgtactgtacag | *Y. lipolytica* gDNA | |
| TGL4up-R | gtcagctccttctagtcttcgatgagtcgtttg | | |
| pGSY1-1F | tctgctggactggatctcggcagtctctcg | pYL-rDNA-CgKCS | pYL-GSY1-CgKCS |
| pGSY1-1R | gagtacagtagtcagagcagattgtactgagag | | |
| Gsup-F | caatctgctctgactactgtactctgaagatc | *Y. lipolytica* gDNA | |
| Gsup-R | taggatgtcctgatactcgtagacggtgac | | |
| pGSY1-2F | gtctacgagtatcaggacatcctactgcgag | pYL-rDNA-CgKCS | |
| pGSY1-2R | gaaaacttgacctgttcggaaatcaacggatg | | |
| Gsdn-F | gatttccgaacaggtcaagttttcggccatg | *Y. lipolytica* gDNA | |
| Gsdn-R | actgccgagatccagtccagcagatcggac | | |
| pSNF1-1F | gaccttcacgttgatctcggcagtctctcg | pYL-rDNA-CgKCS | pYL-SNF1-CgKCS |
| pSNF1-1R | acgtagatatgtcagagcagattgtactgagag | | |
| NFup-F | caatctgctctgacatatctacgtcaatctgac | *Y. lipolytica* gDNA | |
| NFup-R | taggatgtcctgatcggtcataatgttggag | | |
| pSNF1-2F | attatgaccgatcaggacatcctactgcgag | pYL-rDNA-CgKCS | |
| pSNF1-2R | tgaagaggttggtgttcggaaatcaacggatg | | |
| NFdn-F | gatttccgaacaccaacctcttcaagaagatc | *Y. lipolytica* gDNA | |
| NFdn-R | actgccgagatcaacgtgaaggtctcgttg | | |
| pMLC-12R | catcctactgcgcgagtttctgcagcatatcgtcg | pYL-3gD17 | pYL-D17-gMaolE2 |
| GPD-F | tgcagaaactcgcgcagtaggatgtcctgc | | |
| GPD-R | gggggtggccattgttgatgtgtgtttaattcaagaatg | *Y. lipolytica* gDNA | |
| OLE-F | cacacatcaacaatggccaccccccctgcct | | |
| pMLC-1F | ggccgatcttaaccccacgttgccggtctt | pYL-3gD17 | |
| pMLC-1R | gcctccgcctccgcctccgccgcctccgccctcctccttagagtggtcgc | | |
| pMLC-2F | ggcggaggcggcggaggcggaggcggaggctacatcgtgaccagaccc | *Y. lipolytica* gDNA | |
| pMLC-2R | ggcaacgtggggttaagatcggccgttttg | | |
| pPEX10-1F | atttccgaatgggatctcggcagtctctcg | pYL-rDNA-CgKCS | pYL-PEX10-CgKCS |
| pPEX10-1R | cacgtgatcttccagagcagattgtactgagag | | |
| X10up-F | caatctgctctggaagatcacgtggaagaag | *Y. lipolytica* gDNA | |
| X10up-R | taggatgtcctgttgtcgctcatgttggac | | |

| Name | Sequence (5' to 3') | PCR template | Target plasmid |
|---|---|---|---|
| pPEX10-2F | catgagcgacaacaggacatcctactgcgag | pYL-rDNA-CgKCS | |
| pPEX10-2R | tctggttggctctgttcggaaatcaacggatg | | |
| X10dn-F | gatttccgaacagagccaaccagaaagacc | *Y. lipolytica* gDNA | |
| X10dn-R | actgccgagatcccattcggaaatacagtcc | | |
| PEX10-INO2-f | tcgacgataaccccacgttgccggtctt | pYL-PEX10-CgKCS | pYL-PEX10-YlINO2 |
| PEX10-INO2-r | gcagctgcatctgcggttagtactgcaaaaagtgc | | |
| PEX10-INO2-F | ctaaccgcagatgcagctgcgaatcctg | *Y. lipolytica* gDNA | |
| PEX10-INO2-R | caacgtggggttatcgtcgagagcaacatc | | |
| PEX10-INO4-f | tgatgagtaaccccacgttgccggtctt | pYL-rDNA-CgKCS | pYL-PEX10-YlINO4 |
| PEX10-INO4-r | aaaggtgcatctgcggttagtactgcaaaaagtgc | | |
| PEX10-INO4-F | ctaaccgcagatgcacctttcccacccac | *Y. lipolytica* gDNA | |
| PEX10-INO4-R | caacgtggggttactcatcaatcttgggaggatg | | |
| PEX10-35-f | gatcgactaaccccacgttgccggtctt | pYL-PEX10-CgKCS | pYL-PEX10-MoDGAT2-35 |
| PEX10-35-r | agccgtccatctgcggttagtactgcaaaaagtgc | | |
| PEX10-35-F | ctaaccgcagatggacggctcttctcag | Synthesized (Maole_010035) | |
| PEX10-35-R | caacgtggggttagtcgatcttcacgaactc | | |
| PEX10-49-f | aatcatttaaccccacgttgccggtctt | pYL-PEX10-CgKCS | pYL-PEX10-MoDGAT2-49 |
| PEX10-49-r | tctcggccatctgcggttagtactgcaaaaagtgc | | |
| PEX10-49-F | ctaaccgcagatggccgagactgaagtg | Synthesized (Maole_015949) | |
| PEX10-49-R | caacgtggggttaaatgattttcagttctctatcgg | | |
| PEX10-88-f | ctggcagtaaccccacgttgccggtctt | pYL-PEX10-CgKCS | pYL-PEX10-MoGPAT-88 |
| PEX10-88-r | cgaagtccatctgcggttagtactgcaaaaagtgc | | |
| PEX10-88-F | ctaaccgcagatggacttcgtgcccgac | Synthesized (Maole_006088) | |
| PEX10-88-R | caacgtggggttactgccagggctgagac | | |
| PEX10-90-f | tcgacgataaccccacgttgccggtctt | pYL-PEX10-CgKCS | pYL-PEX10-MoGPAT-90 |
| PEX10-90-r | gcagctgcatctgcggttagtactgcaaaaagtgc | | |
| PEX10-90-F | ctaaccgcagatgcagctgcgaatcctg | Synthesized (Maole_006090) | |
| PEX10-90-R | caacgtggggttatcgtcgagagcaacatc | | |
| PEX10-INO2-R | caacgtggggttatcgtcgagagcaacatc | | |

**Table S4.** The amino acid mutations around the binding pocket of AtADS2

| **Mutant** | **Mutation site** |
|---|---|
| AtADS2-M1 | G113A, D117F, A210V, E213V, V214Y, C218V |
| AtADS2-M2 | G113A, A210V, E213V, V214Y, C218V |
| AtADS2-M3 | R37Q, R38V, G113A, D117F, A210V, C218V |
| AtADS2-M4 | R37Q, R38V, G113A, D117F, A210V, E213V, V214Y, C218V |

**Table S5**. The C16 FAEs and C18 fatty acid desaturases evaluated in substrate specificity in this study

| Enzyme | Source | Gene No. | Reference |
|---|---|---|---|
| **C16 FAEs** | | | |
| rELO2 | *Rattus norvegicus* | NM_134383.2 | (Yazawa et al., 2011) |
| CpLCE | *Cryptosporidium parvum* | AAO34582 | (Frltzler et al., 2007) |
| gELOVL6 | *Capra hircus* | NM_001314257.1 | (Shi et al., 2017) |
| **C18 desaturase** | | | |
| D9DMB | *Cunninghamella echinulata* | JN873145 | (Wan et al., 2013) |
| CeFAT6 | *Caenorhabditis elegans* | NM_001268666.1 | (Watts and Browse, 2000) |
| MaOLE2 | *Mortierella alpine* | Y18554.1 | (Wongwathanarat et al., 1999) |

**Table S6**. Concentration of the media used in central composite design in the first-round experiment

| **Sources** | | **Lower Limit** | **Low -1** | **Center 0** | **High +1** | **Higher Limit** |
|---|---|---|---|---|---|---|
| Carbon | Glucose (g/L) | 46.36 | 60 | 80 | 100 | 113.64 |
| Nitrogen | Yeast extract (g/L) | 1.59 | 5 | 10 | 15 | 18.41 |
| | Ammonium sulfate (g/L) | -3.41 | 0 | 5 | 10 | 13.41 |

| **Number** | **Glucose (g/L)** | **Yeast extract (g/L)** | **Ammonium sulfate (g/L)** |
|---|---|---|---|
| 1 | 60 | 5 | 0 |
| 2 | 100 | 5 | 0 |
| 3 | 60 | 15 | 0 |
| 4 | 100 | 15 | 0 |
| 5 | 60 | 5 | 10 |
| 6 | 100 | 5 | 10 |
| 7 | 60 | 15 | 10 |
| 8 | 100 | 15 | 10 |
| 9 | 46.36 | 10 | 5 |
| 10 | 113.64 | 10 | 5 |
| 11 | 80 | 1.59 | 5 |
| 12 | 80 | 18.41 | 5 |
| 13 | 80 | 10 | 0 |
| 14 | 80 | 10 | 13.41 |
| 15 | 80 | 10 | 5 |

**Table S7**. Concentration of the media used in central composite design in the second-round experiment

| Sources | | Lower Limit | Low -1 | Center 0 | High +1 | Higher Limit |
|---|---|---|---|---|---|---|
| Carbon | Glucose (g/L) | 82.96 | 100 | 125 | 150 | 167.05 |
| Nitrogen | Yeast extract (g/L) | 0.64 | 2 | 4 | 6 | 7.37 |
| | Ammonium sulfate (g/L) | 6.64 | 8 | 10 | 12 | 13.36 |

| Number | Glucose (g/L) | Yeast extract (g/L) | Ammonium sulfate (g/L) |
|---|---|---|---|
| 1 | 100.00 | 2.00 | 8.00 |
| 2 | 150.00 | 2.00 | 8.00 |
| 3 | 100.00 | 6.00 | 8.00 |
| 4 | 150.00 | 6.00 | 8.00 |
| 5 | 100.00 | 2.00 | 12.00 |
| 6 | 150.00 | 2.00 | 12.00 |
| 7 | 100.00 | 6.00 | 12.00 |
| 8 | 150.00 | 6.00 | 12.00 |
| 9 | 82.96 | 4.00 | 10.00 |
| 10 | 167.05 | 4.00 | 10.00 |
| 11 | 125.00 | 0.64 | 10.00 |
| 12 | 125.00 | 7.36 | 10.00 |
| 13 | 125.00 | 4.00 | 6.64 |
| 14 | 125.00 | 4.00 | 13.36 |
| 15 | 125.00 | 4.00 | 10.00 |

#### Supplementary Figures

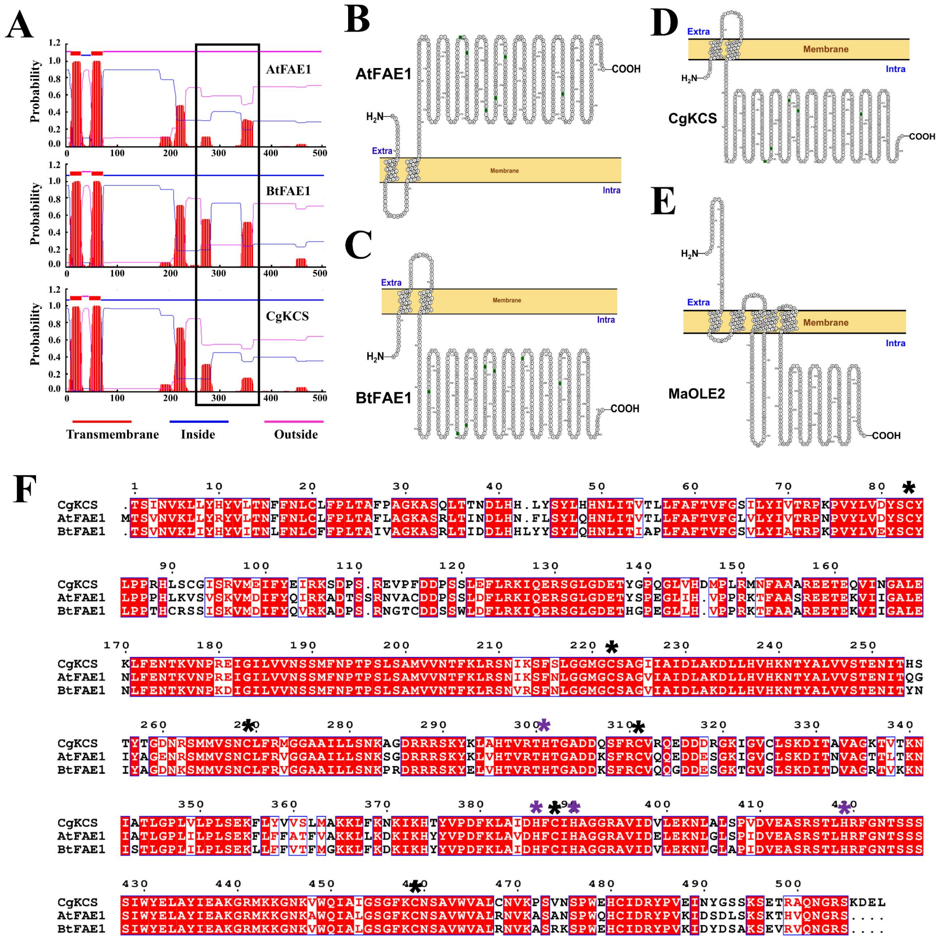


**Fig. S1.** Prediction of transmembrane helices in the fatty acid elongases AtFAE1, BtFAE1, CgKCS and MaOLE2. (A) The pictures showed similar structures among them although they showed distinct capacities on elongating fatty acids in *Y. lipolytica*. The software TMHMM Server v. 2.0 was used. Two-dimensional topology of the entire sequences of AtFAE1 (B), BtFAE1 (C), CgKCS (D) and MaoLE2 (E) constructed by using the online software PROTTER. The catalytic domain of AtFAE1 located in cytosol, while that of the BtFAE1, CgKCS and MaOLE2located in the lumen of ER.

(F) Multiple-sequence alignment of fatty acid elongases. CgKCS, AtFAE1 and BtFAE1 have identical conserved cysteines (black asterisk) and histidine residues (purple asterisk).

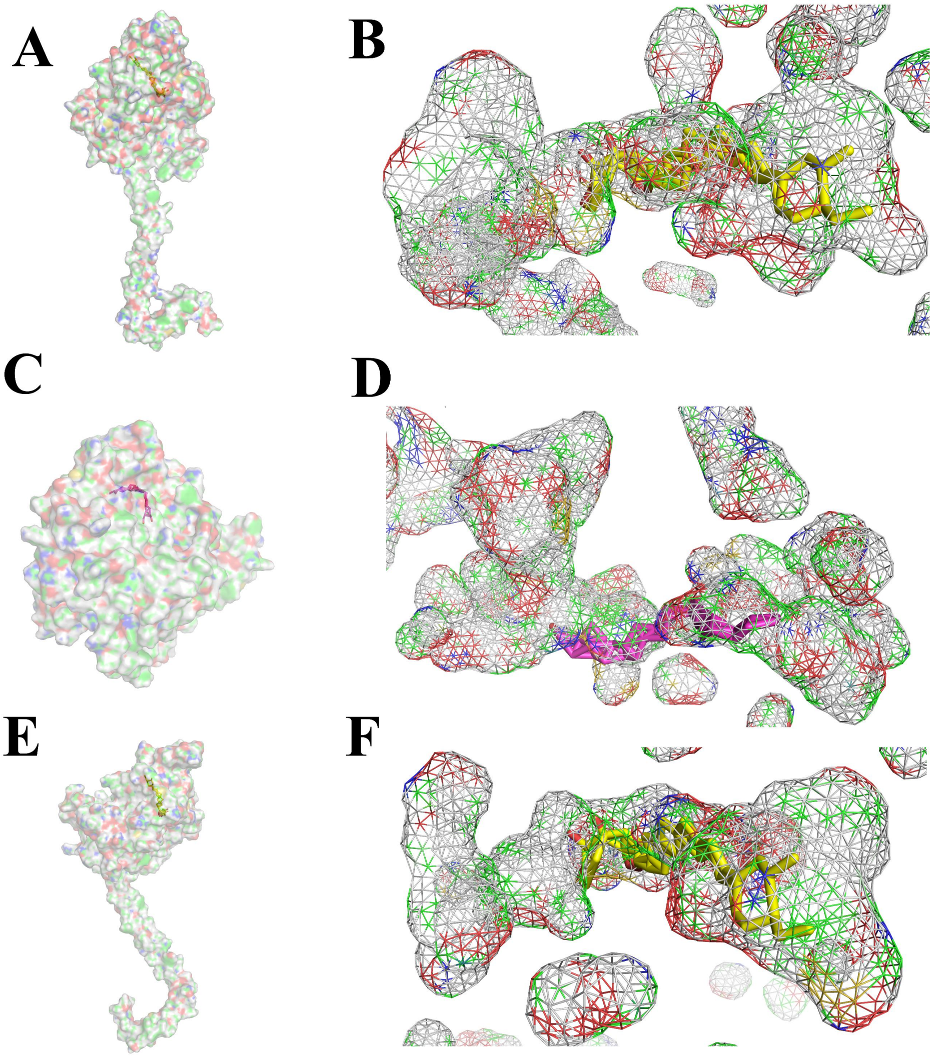


**Fig. S2**. Three-dimensional architectures showing the substrate binding domains in AtFAE1 (A, B), BtFAE1 (C, D) and CgKCS (E, F). The yellow and purple sticks in the catalytic channels denote stearic acid and arachidonic acid, respectively. The enzyme-substrate complexes were modeled according to the crystal structures of *Mycobacterium tuberculosis* PKS11 (PDB ID: 4JAP) for AtFAE1, *Ectocarpus siliculosus* (PDB ID: 4B0N) PKS-I for BtFAE1, and *Mycobacterium tuberculosis* PKS11 (PDB ID: 4JAP) for CgKCS.

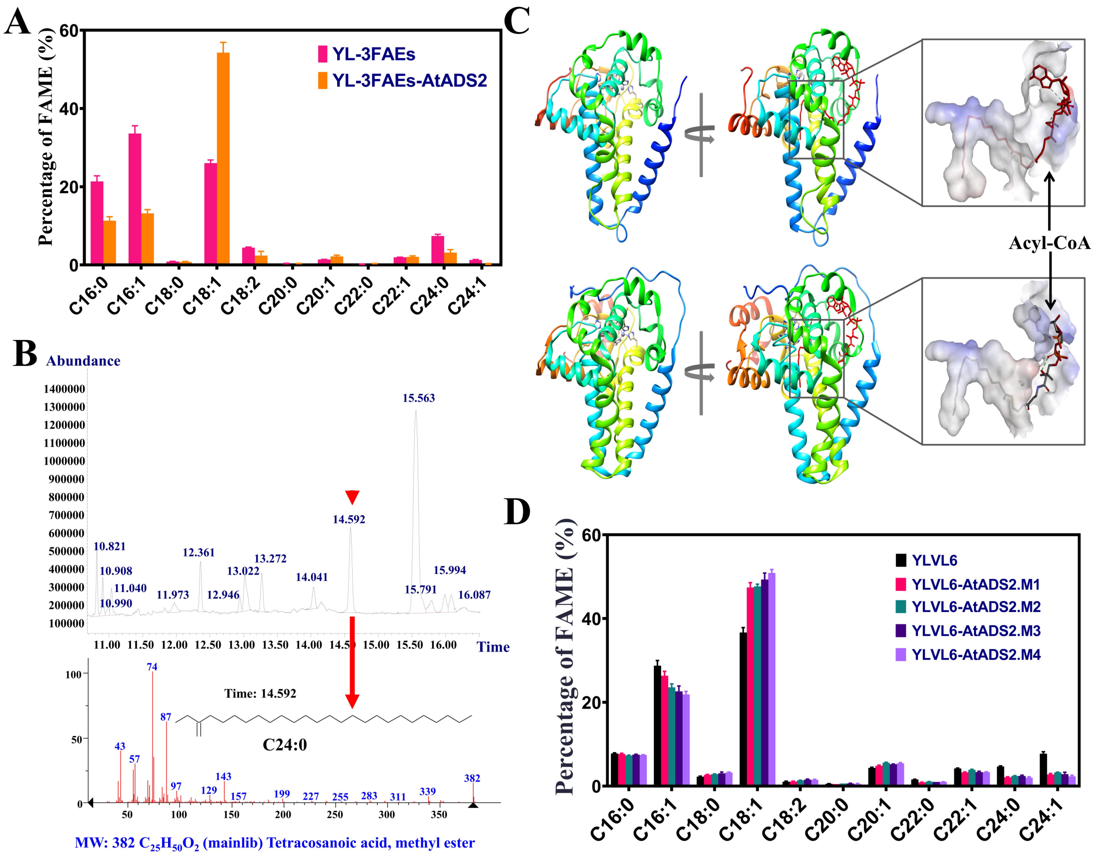


**Fig. S3.** Expression of the acyl-coA desaturase-like protein encoding gene *AtADS2* in *Y. lipolytica*. (A) The production of oleic acid was significantly increased by expressing *AtADS2*. (B) GC-MS detection showed that C24:0-CoA was not desaturated to form C24:1-CoA. (C) Three-dimensional architecture of the C18-acyl-CoA binding domain. The up and down pictures denote AtADS2 and mouse SCD, respectively. The model of the AtADS2/C18-acyl-CoA complex was created by superimposing AtADS2 to the mouse SCD (PDB ID: 4ymk). (D) Expression of *AtADS2* mutants in the strain YLVL6 significantly increased the production of oleic acid but decreased the content of nervonic acid in total fatty acids (TFA).

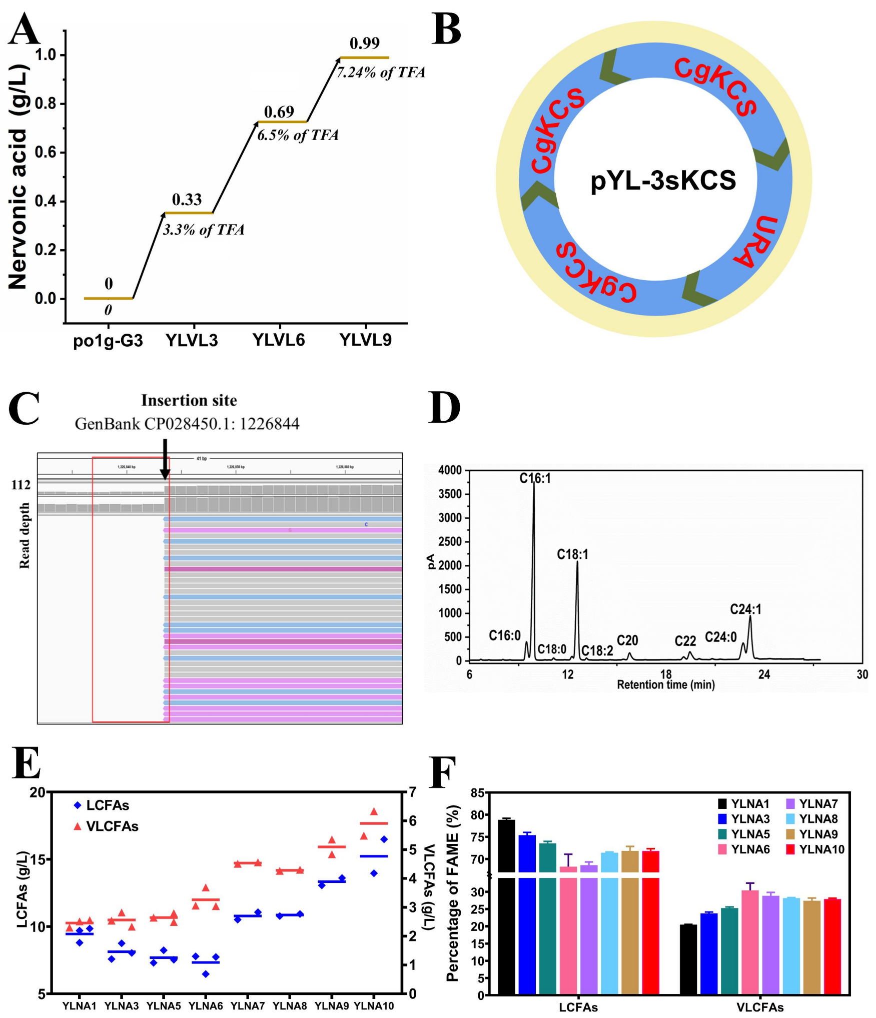


**Fig. S4**. Increasing nervonic acid production by random genetic insertion and homologous recombination. (A) Nervonic acid production increased by iterative expression of *CgKCS*. (B) The arrangement of three copies of CgKCS in the plasmid YL-3sKCS. (C) Identifying the insertion sites of the *CgKCS* expression cassettes by genome resequencing. The picture was generated by the Integrative Genomics Viewer (IGV) software. The insertion site CP028450.1: 1226844 located in the elongation factor 1-alpha (TEF1) gene in *Y. lipolytica*. The results of lipid and VLCFAs biosynthesis in the strains YLNA1, 3, and 5 to 10, including the fatty acid composition in YLNA9 (D), the titer of LCFAs and VLCFAs (E), and the fraction of LCFAs and VLCFAs (F). Data are mean ± s.d. from three replicates.

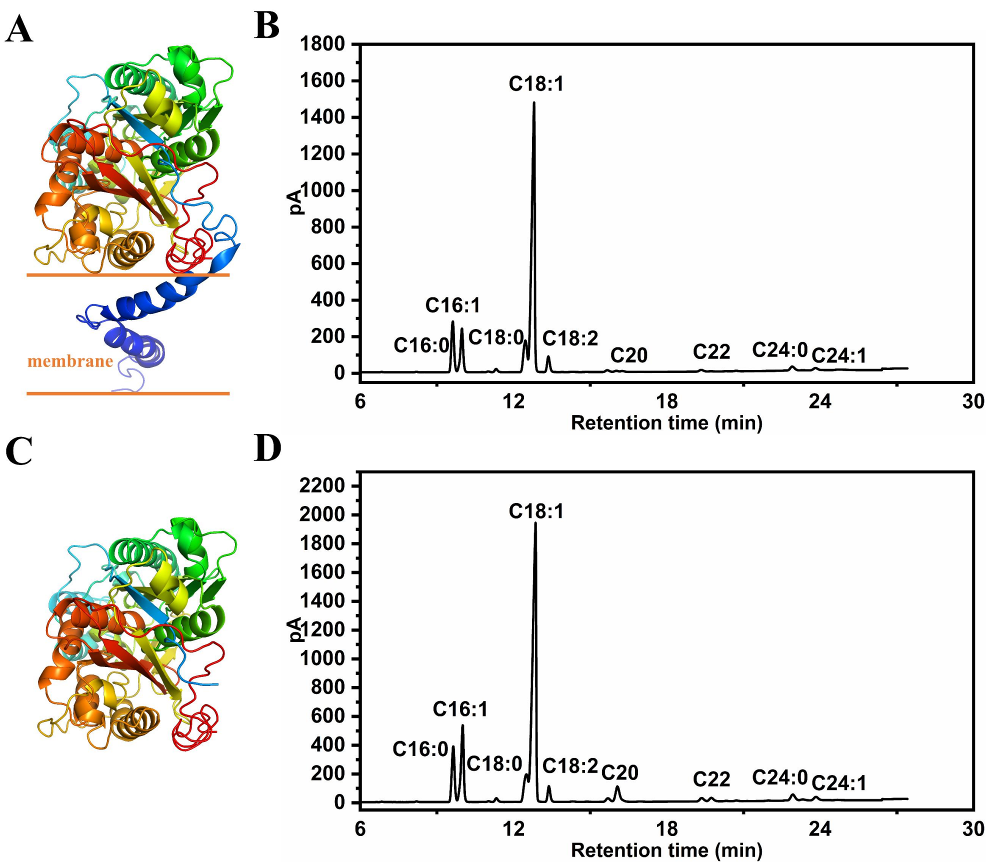


**Fig. S5**. The influence of the transmembrane region on the catalytic ability of CgKCS. The structural models of CgKCS (A) and the truncated CgKCS (C). The fraction of fatty acids in the control strain (B) and the truncated CgKCS expressing strain (D).

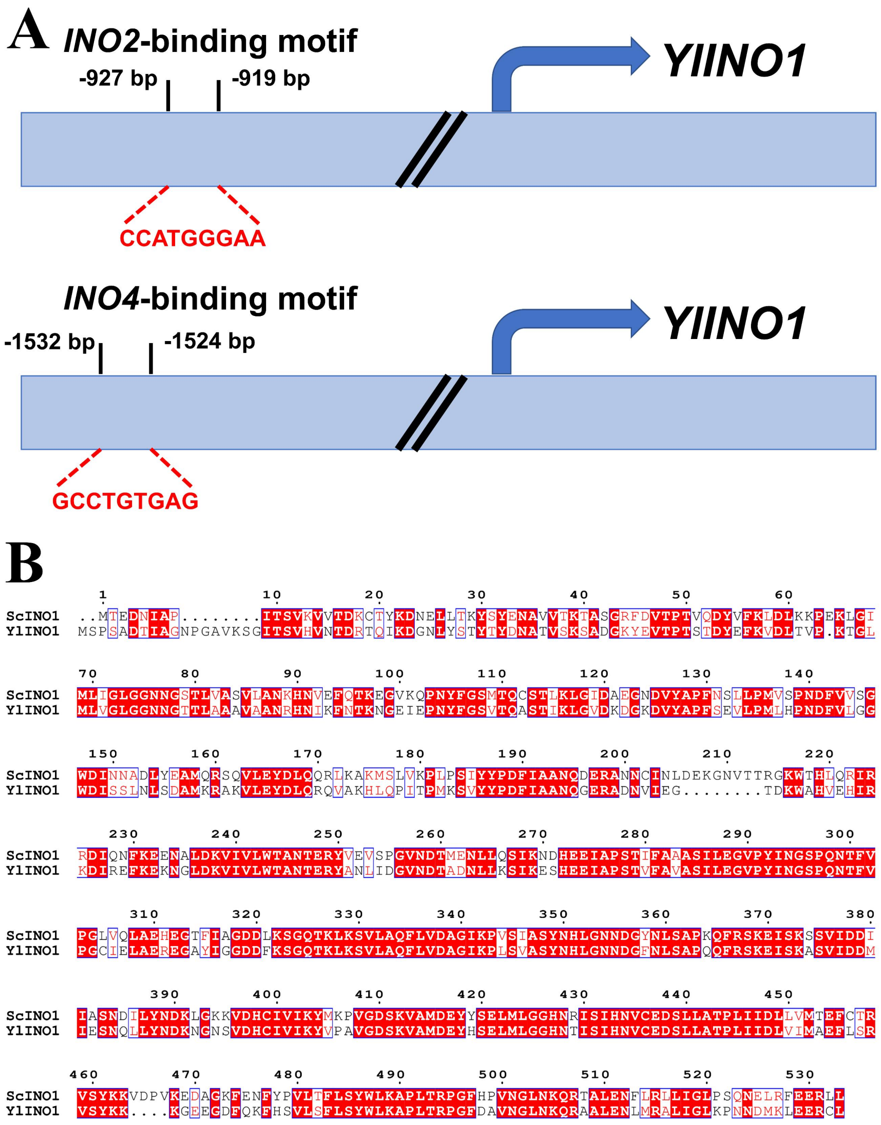


**Fig. S6**. Predicting the interaction of *S. cerevisiae* INO2/INO4 with the promoter of the *Y. lipolytica INO1*. (A) Schematic diagram showing the ScINO2 and ScINO4 binding motifs in the YlINO1 promoter region. (B) Multiple-sequence alignment of INO1 proteins. Sc, *Saccharomyces cerevisiae*; Yl, *Yarrowia lipolytica*. Conserved amino acids are denoted in red boxes.

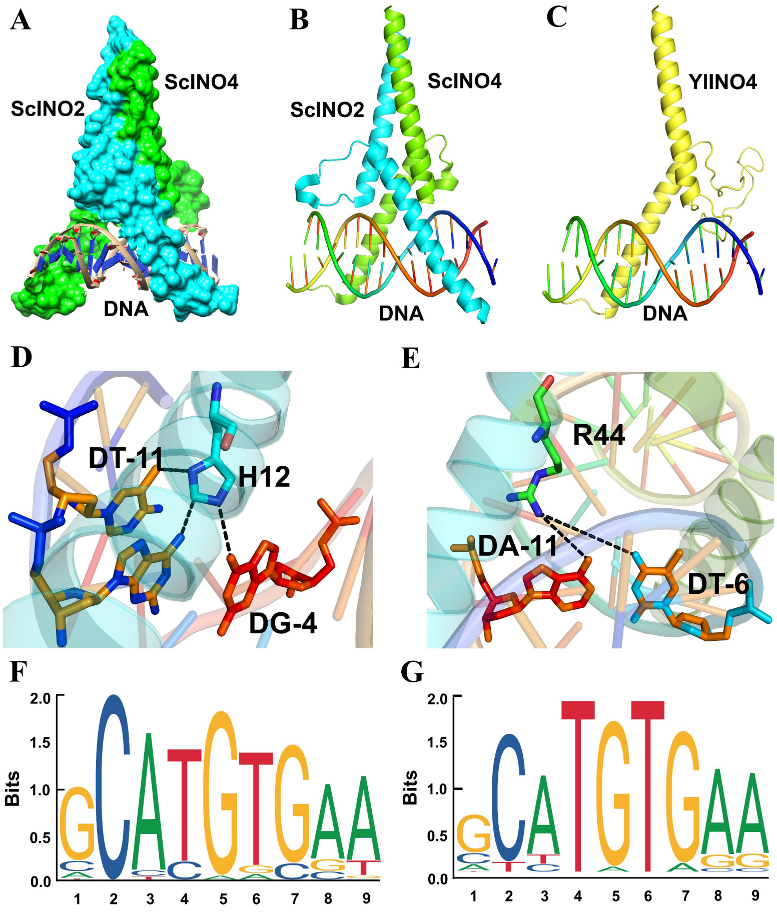


**Fig. S7.** Three-dimensional architecture and sequence frequency analysis of ScINO2/ScINO4. (A) and (B) Overall three-dimensional structural model of the surface of INO2/INO4 complex. (C) Three-dimensional architecture of the complex of YlINO4 combining with DNA. (D) and (E) The interaction between ScINO2 and the promoter. Black dash lines denote the interaction between critical amino acid residues and nucleic acids. The sequence frequency of ScINO2 (F) and ScINO4 (G).

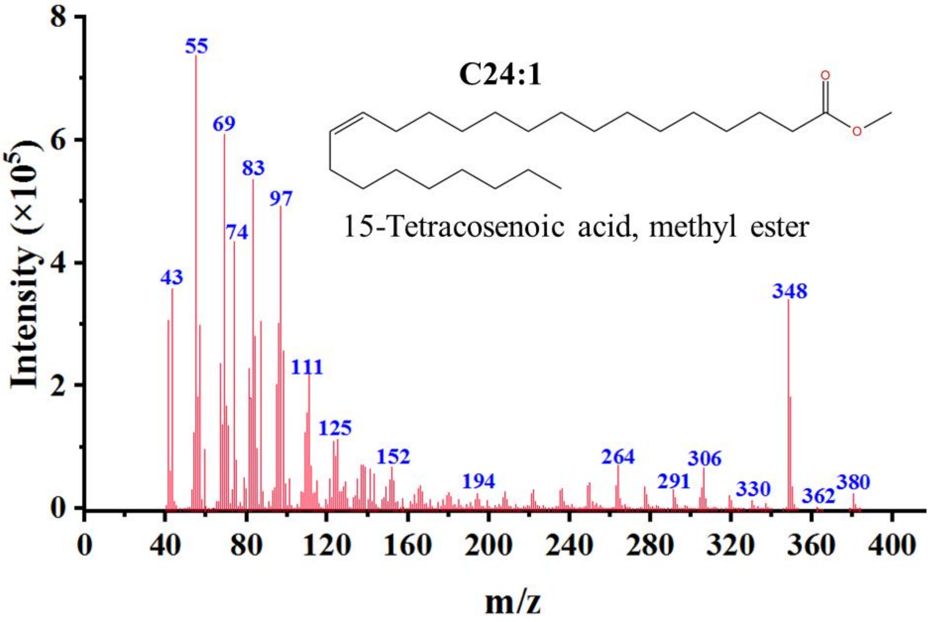


**Fig. S8.** Identification of nervonic acid in the lipids produced by the engineered *Y. lipolytica* strains by GC-MS.

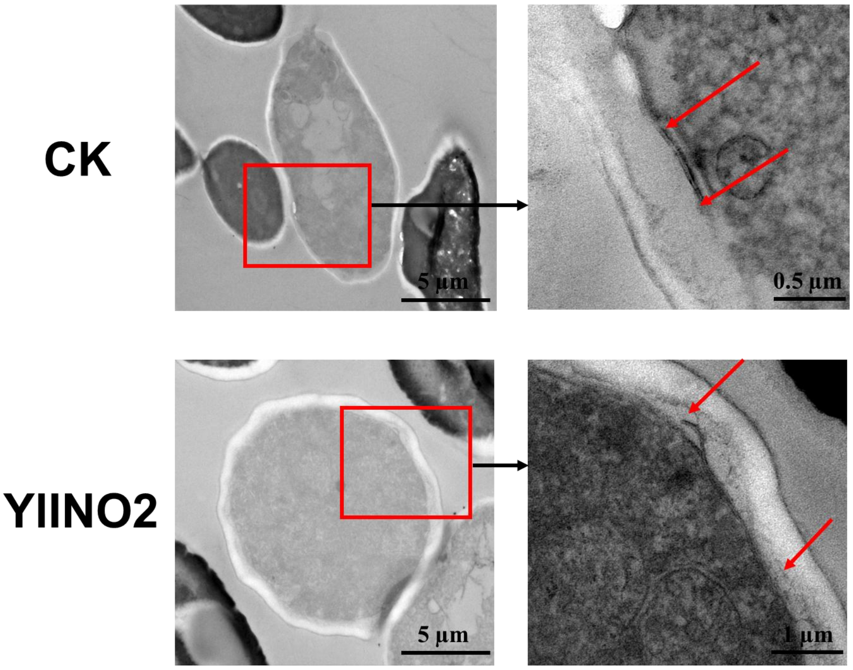


**Fig. S9**. Overexpression of YlINO2 enlarged the size of ER. Transmission electron micrographs of the CK (YLNA8) and YlINO2 (YLNA9) strains. The red arrows indicate the two ends of ER.

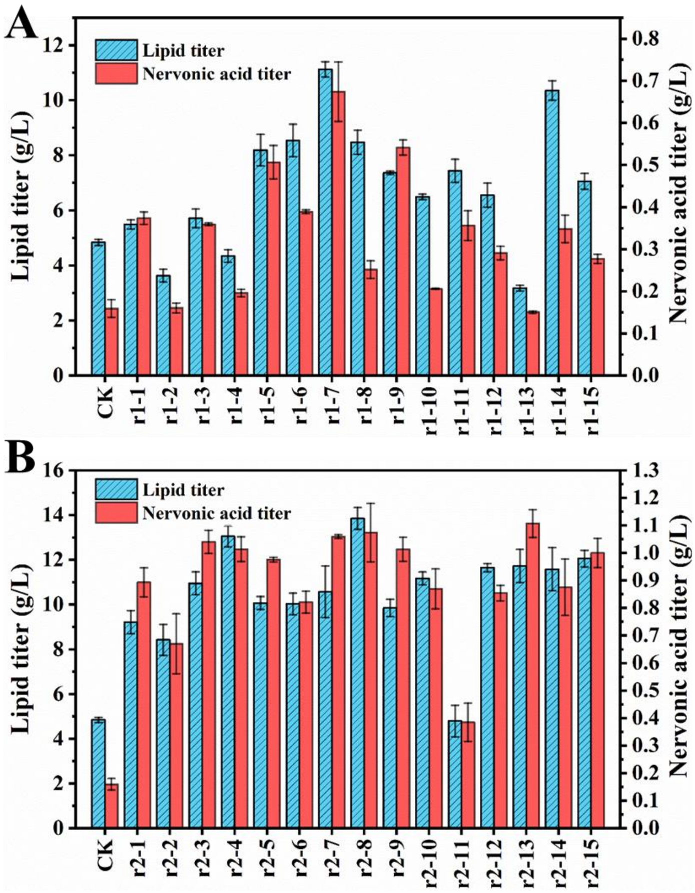


**Fig. S10.** Lipid and nervonic acid production by the strain YLNA3 in different media designed by the central composite design (CCD) and response surface methodology. (A) The first round of medium optimization. The medium r1-7 was used in next optimization. (B) The second round of medium optimization. The medium r2-8 was used in the following fermentation.

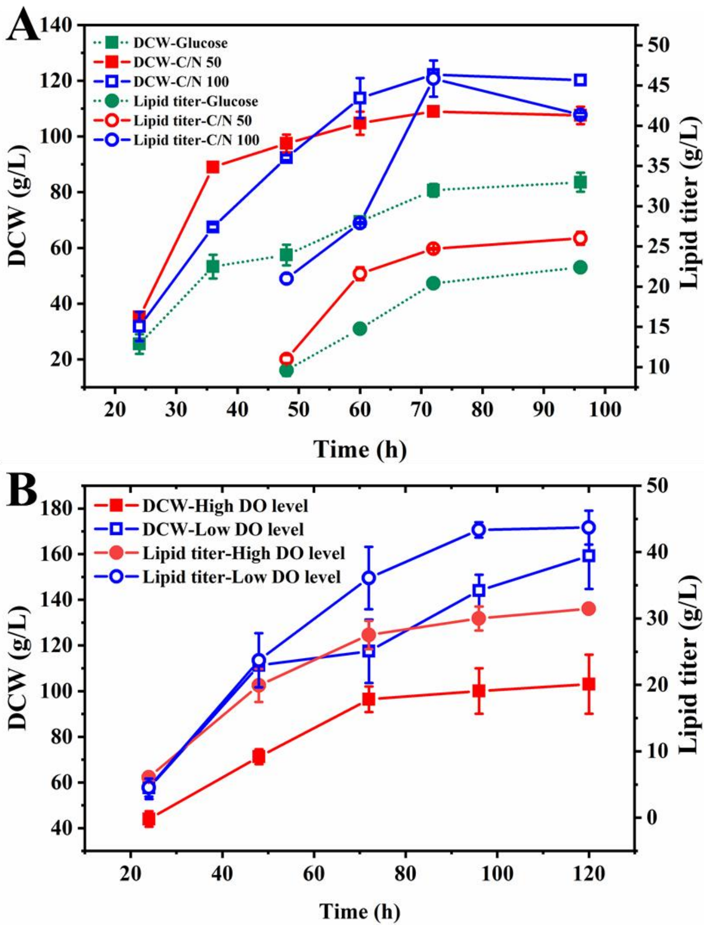


**Fig. S11.** Optimization of feeding medium composition (A) and dissolved oxygen level (B) in a 3-L reactor. Feeding glucose and ammonium sulfate with a C/N molar ratio of 100:1 during fermentation resulted in an 84.6% increase in lipid titer compared to feeding sole glucose (A). The lipid titer was further improved to 43.7 g/L by controlling the dissolved oxygen level at 20% in 24 h and below 5% after 24 h (B). Data are mean ± s.d. from two replicates.

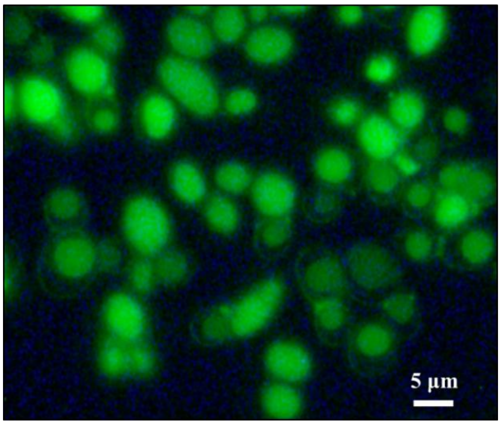


**Fig. S12**. Observation of the lipid accumulation in the strain YLNA9 by Nile red dyeing. The cells cultivated for 216 h in the 50-L reactor was used. The green color represents the lipid droplets in cells.

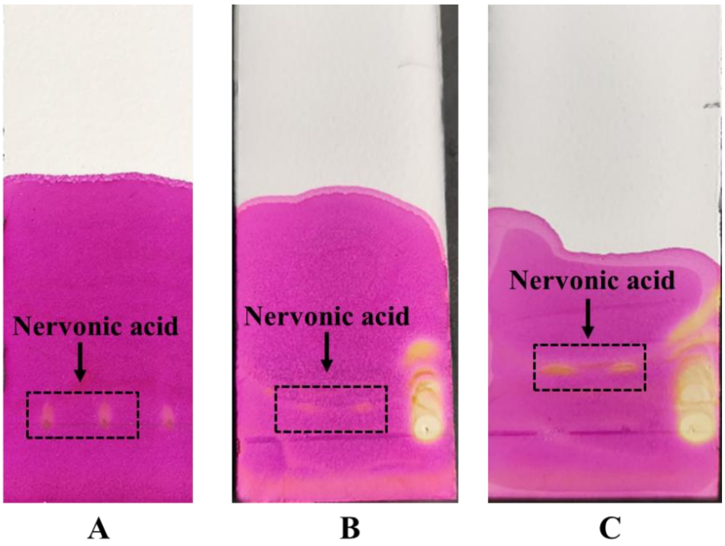


**Fig. S13**. Evaluation of the elution buffers used in silica gel column chromatography for nervonic acid separation by thin layer chromatography (TLC). The developing buffers included *n*-hexane supplemented with 0.5% ammonia (A), 0.5% acetic acid (B), and 1% acetic acid (C). The outcome showed that the buffer consisting of *n*-hexane and 1% acetic acid exhibited best separation efficiency and was used in silica gel column chromatography.

**Reference**

Frltzler, J.M., Millership, J.J., Zhu, G., 2007. *Cryptosporidium parvum* long-chain fatty acid elongase. Eukary. Cell. 6, 2018-2028.

Li, J.X., Xu, J., Ruan, J.C., Meng, H.M., Su, H., Han, X.F., et al., 2020. Disrupting a phospholipase A2 gene increasing lipid accumulation in the oleaginous yeast *Yarrowia lipolytica*. J. Appl. Microbiol. 130, 100-108.

Shi, H.B., Wu, M., Zhu, J.J., Zhang, C.H., Yao, D.W., Luo, J., Loor, J.J., 2017. Fatty acid elongase 6 plays a role in the synthesis of long-chain fatty acids in goat mammary epithelial cells. J. Dairy. Sci. 100, 4987-4995.

Wan, X., Liang, Z., Gong, Y., Zhang, Y., Jiang, M., 2013. Characterization of three Delta9-fatty acid desaturases with distinct substrate specificity from an oleaginous fungus *Cunninghamella echinulata*. Mol. Biol. Rep. 40, 4483-4489.

Watts, J.L., Browse, J., 2000. A palmitoyl-CoA-specific Δ9 fatty acid desaturase from *Caenorhabditis elegans*. Biochem. Biophys. Res. Commun. 272, 263-269.

Wongwathanarat, P., Michaelson, L.V., Carter, A.T., Lazarus, C.M., Griffiths, G., Stobart, A.K., et al., 1999. Two fatty acid Δ9-desaturase genes, *ole1* and *ole2*, from *Mortierella alpina* complement the yeast *ole1* mutation. Microbiol-SGM. 145, 2939-2946.

Yazawa, H., Kamisaka, Y., Kimura, K., Yamaoka, M., Uemura, H., 2011. Efficient accumulation of oleic acid in *Saccharomyces cerevisiae* caused by expression of rat elongase 2 gene (*rELO2*) and its contribution to tolerance to alcohols. Appl. Microbiol. Biot. 91, 1593-1600.
